## Supplemental Material for "Idebenone Enhances the Early Microglial Response to Traumatic Brain Injury and Mitigates Acute Gene Expression Changes to Ephrin-A and Dopamine Signaling Pathways"

**Supplemental Figure S1.** Outlier-based Npas4 exclusion

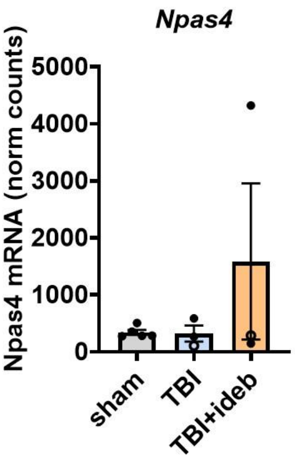

**Supplemental Table 3.** Complete list of genes differentially expressed between the TBI+vehicle and TBI+idebenone groups with p-values between 0.05 and 0.1. The genes shown in bold encode proteins reported to interact with the idebenone-binding protein SHC1, or with SHC1-interacting receptors. The underlined genes are preferentially expressed by microglia within the healthy mouse brain [19].

| Name | Description | Fold Change | p-Value |
|---|---|---|---|
| Adora2a | adenosine A2a receptor | 4.23 | 0.057 |
| **Ntrk1** | neurotrophic tyrosine kinase, receptor, type 1 | 2.12 | 0.062 |
| Adcy5 | adenylate cyclase 5 | 1.82 | 0.057 |
| Pde1b | phosphodiesterase 1B, Ca2+-calmodulin dependent | 1.66 | 0.068 |
| Gabra4 | gamma-aminobutyric acid (GABA) A receptor, subunit alpha 4 | 1.29 | 0.067 |
| <u>Irf8</u> | interferon regulatory factor 8 | 1.28 | 0.076 |
| Ccnd1 | cyclin D1 | 1.28 | 0.095 |
| Nol3 | nucleolar protein 3 (apoptosis repressor with CARD domain) | 1.27 | 0.075 |
| <u>Nfkbia</u> | nuclear factor of kappa light polypeptide gene enhancer in B cells inhibitor, alpha | 1.23 | 0.096 |
| <u>C1qc</u> | complement component 1, q subcomponent, C chain | 1.23 | 0.050 |
| <u>P2rx4</u> | purinergic receptor P2X, ligand-gated ion channel 4 | 1.22 | 0.079 |
| Kcna1 | potassium voltage-gated channel, shaker-related subfamily, member 1 | 1.20 | 0.086 |
| S100b | S100 protein, beta polypeptide, neural | 1.19 | 0.081 |
| Eng | endoglin | 1.18 | 0.088 |
| Ppp3ca | protein phosphatase 3, catalytic subunit, alpha isoform | 1.18 | 0.059 |
| <u>Cx3cr1</u> | chemokine (C-X3-C motif) receptor 1 | 1.15 | 0.070 |
| Gnb5 | guanine nucleotide binding protein (G protein), beta 5 | 1.14 | 0.088 |
| **Igf1r** | insulin-like growth factor I receptor | 1.13 | 0.061 |
| Cntnap2 | contactin associated protein-like 2 | 1.13 | 0.093 |
| Nfe2l2 | nuclear factor, erythroid derived 2, like 2 | 1.13 | 0.085 |
| Mapt | microtubule-associated protein tau | 1.12 | 0.072 |
| Glrb | glycine receptor, beta subunit | 1.12 | 0.055 |
| **Src** | Rous sarcoma oncogene | 1.12 | 0.099 |
| Chd4 | chromodomain helicase DNA binding protein 4 | 1.11 | 0.090 |
| **Rhoa** | ras homolog gene family, member A | 1.11 | 0.060 |
| **App** | amyloid beta (A4) precursor protein | 1.07 | 0.058 |
| Dld | dihydrolipoamide dehydrogenase | -1.06 | 0.098 |
| Nell2 | NEL-like 2 | -1.07 | 0.072 |
| Il6 | interleukin 6 | -1.25 | 0.094 |
| Casp6 | caspase 6 | -1.26 | 0.099 |
| **Ngf** | nerve growth factor | -1.50 | 0.072 |
| Epha3 | Eph receptor A3 | -1.71 | 0.078 |

**Supplemental Table 4.** Results of "ENCODE and ChEA Consensus TFs from ChIP-X" Enrichr query for idebenone-affected genes following expansion of the list to include the *Drd2*-correlated genes in Figure 10a-c. The adjusted p-value (p-adj) was calculated using the Benjamini-Hochberg method for correction for multiple hypotheses testing.

| term | p-value | p-adj value | overlap_genes |
|---|---|---|---|
| **SUZ12** ChEA | 3.752209e-12 | 2.101237e-10 | Adcy5, Adora2a, Bcl2, Calb1, Camk4, Cxcl12, Ccxr4, Drd2, Efna5, Epha3, Epha5, Epha6, Epha7, Gabra4, Gad2, Htr1a, Mal, Mmp9, Negr1, Npy, Ntf3 Slc32a1 |
| **ESR1** ChEA | 0.000466 | 0.012189 | Bcl2, Cxcl12, Efna1, Itpr1 |
| **REST** ChEA | 0.000653 | 0.012189 | Adcy5, Calb1, Drd1, Drd2, Fgf14, Gad2, Htr1a, Slc28a3, Slc32a1, Tenm2 |
| **REST** ENCODE | 0.001988 | 0.027833 | Drd2, Calb1, Chat, Grm2, Htr1a |
| **EZH2** ChEA | 0.018165 | 0.203448 | Adcy5, Camk4, Gad2 |
| **AR** ChEA | 0.042033 | 0.392305 | Ang, Efna1 |
